# SpiderSeqR: an R package for crawling the web of high-throughput multi-omic data repositories for data-sets and annotation

**DOI:** 10.1101/2020.04.13.039420

**Authors:** Anna M. Sozanska, Charles Fletcher, Dóra Bihary, Shamith A. Samarajiwa

## Abstract

More than three decades ago, the microarray revolution brought about high-throughput data generation capability to biology and medicine. Subsequently, the emergence of massively parallel sequencing technologies led to many big-data initiatives such as the human genome project and the encyclopedia of DNA elements (ENCODE) project. These, in combination with cheaper, faster massively parallel DNA sequencing capabilities, have democratised multi-omic (genomic, transcriptomic, translatomic and epigenomic) data generation leading to a data deluge in bio-medicine. While some of these data-sets are trapped in inaccessible silos, the vast majority of these data-sets are stored in public data resources and controlled access data repositories, enabling their wider use (or misuse). Currently, most peer reviewed publications require the deposition of the data-set associated with a study under consideration in one of these public data repositories. However, clunky and difficult to use interfaces, subpar or incomplete annotation prevent discovering, searching and filtering of these multi-omic data and hinder their re-purposing in other use cases. In addition, the proliferation of multitude of different data repositories, with partially redundant storage of similar data are yet another obstacle to their continued usefulness. Similarly, interfaces where annotation is spread across multiple web pages, use of accession identifiers with ambiguous and multiple interpretations and lack of good curation make these data-sets difficult to use. We have produced SpiderSeqR, an R package, whose main features include the integration between NCBI GEO and SRA databases, enabling an integrated unified search of SRA and GEO data-sets and associated annotations, conversion between database accessions, as well as convenient filtering of results and saving past queries for future use. All of the above features aim to promote data reuse to facilitate making new discoveries and maximising the potential of existing data-sets.

**Availability:** https://github.com/ss-lab-cancerunit/SpiderSeqR

## 1 INTRODUCTION

In the not so distant past, molecular techniques such as Southern and Northern blotting enabled interrogation of DNA and mRNA molecules, a single DNA fragment or gene at a time. The invention of DNA microarrays in the mid-1990’s [1] enabled the simultaneous measurement of expression of tens of thousands of genes. Commercialization and development of different array platforms for gene expression, single nucleotide polymorphism (SNP) and copy number aberration detection by Agilent, Affymetrix, Illumina and others led to wider uptake of these technologies and a data explosion in bio-medicine. Subsequently, with the development of massively parallel sequencing, generation of cheaper, genome scale data-sets has become a reality. During the past two decades, proliferation of large scale sequencing projects such as the Human Genome Project, 1000 Genomes, UK 100K Genomes, Encyclopedia of DNA elements (Encode) project, Genotype Tissue Expression (GTEx), and Roadmap Epigenomic project have democratised sequencing data generation and usage. with 30X whole genome sequence costing only a few hundred US dollars, it has been estimated that currently, exabyte scale data-sets are being generated by sequencing technologies alone [2]. Whilst sequencing cost is rapidly decreasing, the cost associated with carrying out well designed replicated experiments as well as generating sequencing libraries still make these ‘omic’ data-sets expensive to produce, store and analyse.

A number of distinct sequencing data repositories have been established to store these open access (or controlled access - usually from human subjects) data. The National Centre for Biotechnology Information Sequence Read Archive (NCBI SRA: https://www.ncbi.nlm.nih.gov/sra) [3, 4] as well as controlled access repositories such as the database of genotypes and phenotypes (dbGaP: https://www.ncbi.nlm.nih.gov/gap) [5], the European Genome Phenome Archive (EGA: https://ega-archive.org) [6] are some of these large public sequence data repositories. They store data-sets from multiple sequencing platforms, cell types, species and technologies. Similarly NCBI Gene Expression Omnibus (NCBI GEO: https://www.ncbi.nlm.nih.gov/geo) [7] and ArrayExpress (https://www.ebi.ac.uk/arrayexpress) [8] are repositories that archive high-throughput gene expression and other functional genomic data-sets. The above repositories enable data-sets generated to be re-purposed to ask new scientific questions, generate new hypotheses, making it possible for several studies to be pooled to increase statistical power to produce findings that couldn’t have been achieved from smaller individual studies [9]. This also enables speedier translation of research findings into clinical practice, preventing data being trapped in private silo’s. Access to data-sets from published studies also enhances transparency and reproducibility. Consequently, high quality data increases collaboration, interdisciplinary research and computational methodology development.

Currently, many journals require authors to submit their data to these repositories and provide an accession number prior to publication. Accordingly, funding and governmental bodies in the United States, United Kingdom and the European Union, including the National Institutes of Health (NIH), and the National Science Foundation (NSF), UK Research and Innovation councils and the European Commission, have instituted policies and issued statements in support of data sharing and open science. In order to fully utilise the potential of the public ‘omic’ data repositories, efficient search tools are needed to identify data-sets, associated metadata and samples of interest. A number of such tools are in existence for a variety of platforms. However, searching for large volumes of experiments for specific data-sets still remains a very labour-intensive process. SpiderSeqR is an R package [10] that aims to fulfill this need and provides a modular tool-kit for more efficient access to the NCBI SRA and GEO databases.

## 2 METHODS

### 2.1 Package dependencies

The Bioconductor project [11] is an open-source software development project using the R statistical programming environment [10] for the analysis and comprehension of genomic data. According to one of the key tenets of the Bioconductor project [11], where possible new tools should reuse the functionality of existing methods to avoid reinventing existing functionality. SpiderSeqR relies on the database files generated by Bioconductor’s SRAdb and GEOmetadb packages. Other package dependencies include SQLite interface (DBI, RSQLite) [12] and tidy data functionality provided by (dplyr, plyr and tidyr) [13].

### 2.2 Package overview

SpiderSeqR offers a set of modular and convenient tools for streamlining the process of searching for ‘omic’ metadata and data-sets of interest. Its main functionalities include a number of distinct search methods, accession conversion, modifying and displaying the results and saving and re-running the query in the future. The main advantage of the package is that it integrates data from GEO and SRA databases and provides novel methods for speeding up the process of discovering appropriate sets of samples and associated metadata for answering novel questions using publicly available data-sets.

### 2.3 Database overview

SpiderSeqR integrates and interconnects NCBI SRA and GEO databases; where possible, metadata is gathered from both of these databases. The two databases have different hierarchies for arranging submissions, as shown in Fig. 1. SRA’s top *accession level* is study (SRP), followed by sample (SRS), experiment (SRX) and run (SRR). GEO uses series (GSE) and samples (GSM). However, the structure of GEO is complicated by some of the GSEs taking a role of *superseries*, where one GSE can link other GSEs (and in that case a given GSM can belong to multiple GSEs; something that never happens in SRA). There is some degree of overlap between the two databases. The numbers in the centre of Fig. 1 indicate the count of each accession level with corresponding information in the other database (e.g. there are 1 091 384 GSMs which can be converted to SRA accessions). The GSEs are not included in the overlapping counts because their count is less informative owing to the presence of superseries. On the whole, one can generalise that SRPs correspond to GSEs and GSMs can be converted to SRXs (or SRSs); however, there are some exceptions, so each study/series is considered independently. SpiderSeqR relies on a custom built database for conversion between the accession levels between SRA and GEO.

**Figure 1:**
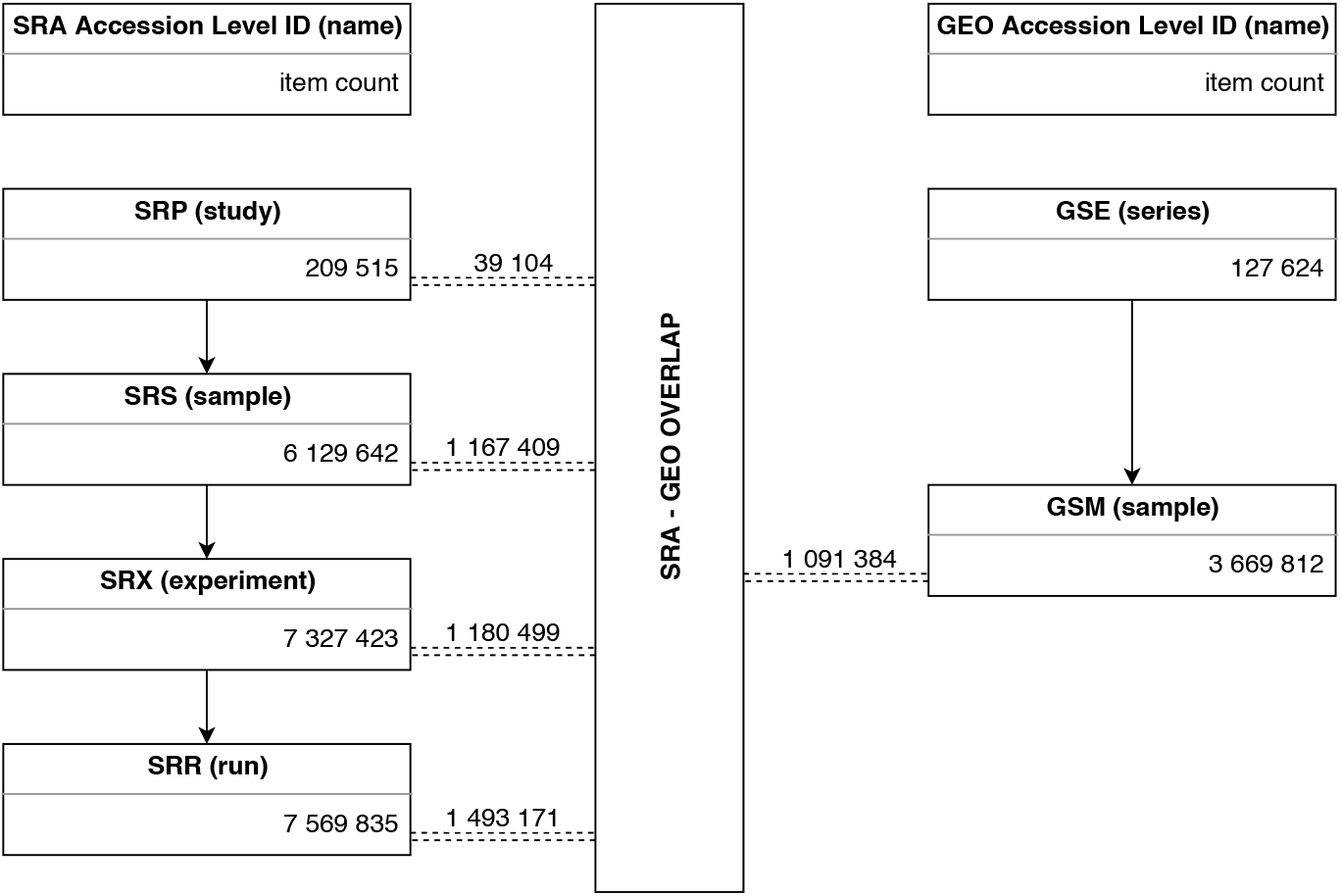
Overview of the hierarchy of accessions within NCBI SRA and GEO databases

### 2.4 Main functions

An overview of the package functions, grouped broadly according to their purpose, is shown in Fig.2. The most important functions are briefly introduced below.

**Figure 2:**
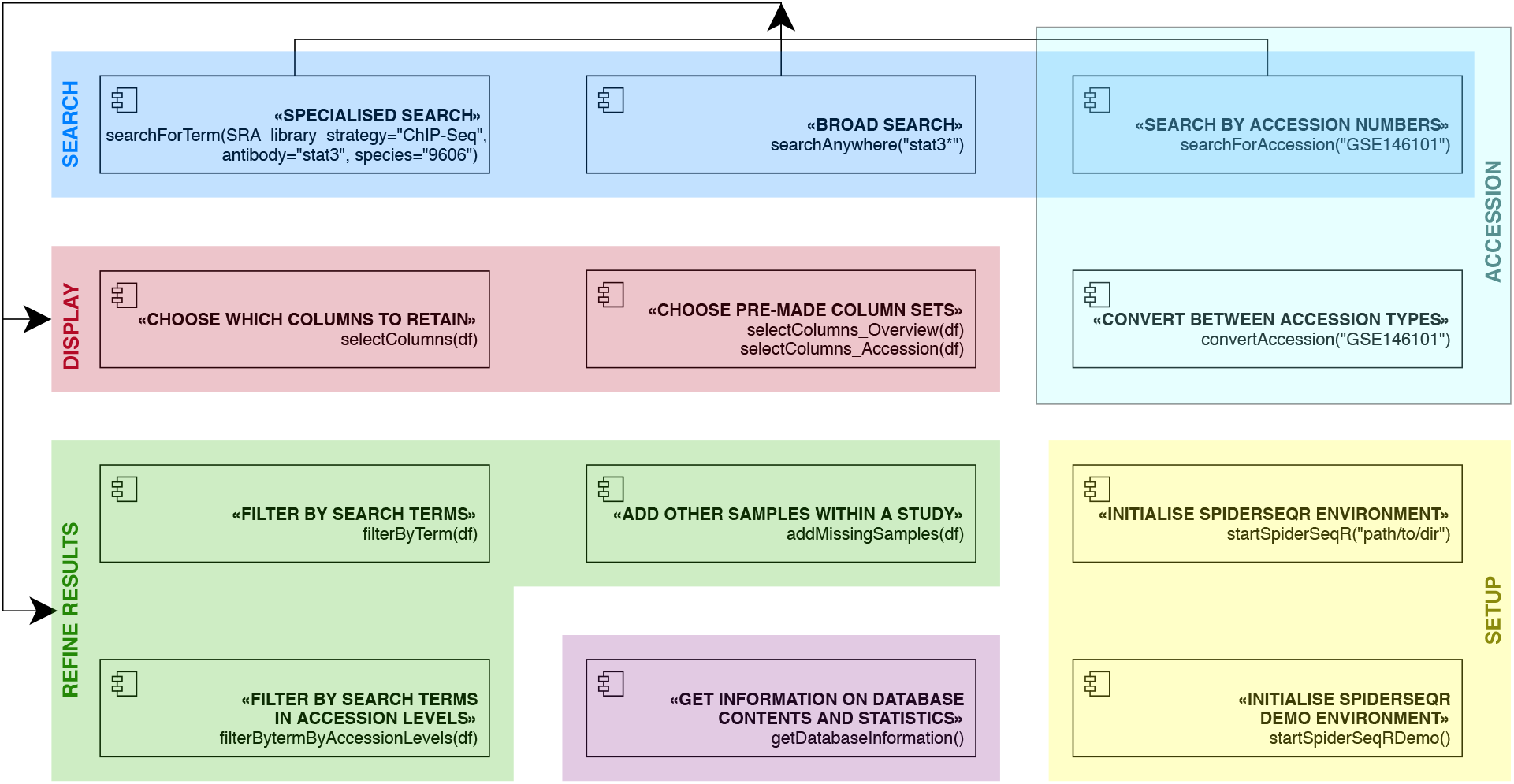
Overview of the functions within SpiderSeqR package

`searchForAccession()` searches for specific accessions and provides all associated meta information, e.g. `searchForAccession(“SRPꙩ1ꙩꙩ68”)`. Any accession type within SRA and GEO can be supplied.

`searchAnywhere()` provides full-text search (by default only within SRA; search within GEO is only possible if the user chooses to create the fulltext search database), e.g. `searchAnywhere(“stat3”)`. This allows for google-like search, which includes all fields that were included in the submission (there is also an option of restricting search to fields corresponding to selected *accession levels*). The use of wildcard ‘*’ is also allowed, e.g. `searchAnywhere(“stat*”)`, which would return any matches to stat1, stat2, etc. (as well as any other letters or numbers such as ‘static’). The main aim of this function is to search for all potential matches, which can be further refined using other SpiderSeqR tools.

`searchForTerm()` is a search function with a custom-designed logic optimised to filter out incorrect matches. It also attempts to label input samples in ChIP-Seq experiments and control samples in RNA-Seq, however, due to inconsistencies of GEO/SRA metadata, we strongly recommend visually checking the output to ensure ChIP-seq input and RNA-seq controls are properly labelled. E.g. `searchForTerm(library_strategy = “ChIP-Seq”, antibody = c(“p53”, “tp53”), species = c(“96ꙩ6”, “1ꙩꙩ9ꙩ”))` searches for ChIP-Seq experiments on human or mouse cells using p53 antibody (synonyms can be supplied within the vector).

`convertAccession()` converts between all available accession types, e.g. `convertAccession(“SRPꙩ1ꙩꙩ68”)`. It provides a data frame with all possible accession levels and their conversion between SRA and GEO (i.e. SRP, SRX, SRS, SRR, GSE, GSM), if available. Just like with `searchForAccession()`, any accession type within SRA and GEO can be supplied.

`filterByTerm()` allows users to filter and refine their search results. This is especially useful for refining the results of queries which produced a large number of matching samples.

`filterByTermByAccession()`, similarly to `filterByTerm()`, allows users to filter and refine their search results. However, only information pertaining to a given set of *accession levels* is used for filtering.

`rerunSpiderSeqR()` allows users to run previous queries again. This ensures reproducible querying and also allows users to run the query again on the updated versions of the database to find new results.

`getDatabaseInformation()` allows users to explore some of the information about the databases; simply use `getDatabaseInformation()` to access an interactive menu with a list of available options (e.g. total number of studies in SRA, most common library strategies, etc.).

## 3 RESULTS

### 3.1 SpiderSeqR Workflow

SpiderSeqR can be used for a variety of search types and purposes. However, one key application is for collection of data-sets, which will be used for a high-throughput analysis in big-data approach to verifying novel biological hypotheses.

For example, one might be interested in finding all ChIP-Seq data-sets where *MYC*-directed antibodies were used to obtain *MYC* binding in various cellular contexts. An example of such a SpiderSeqR workflow is illustrated in Fig. 3. The most convenient way of conducting a query is to use:

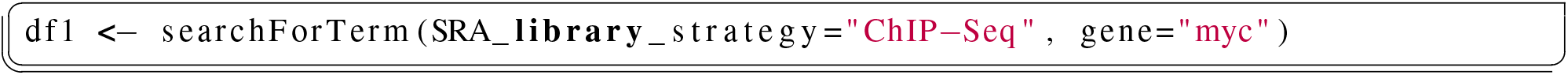

**Figure 3:**
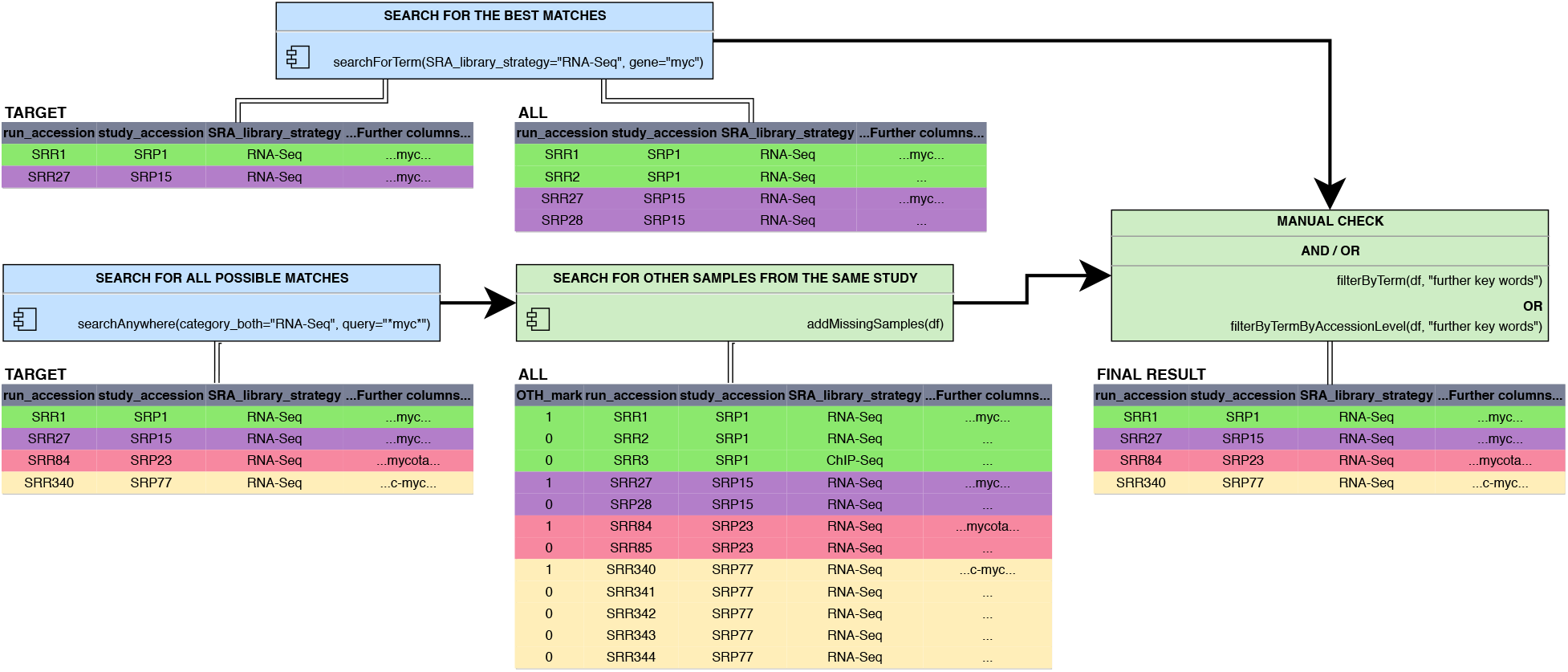
An example of the workflow with SpiderSeqR package

This is a very specific search, but comes at the expense of potentially missing some samples (which are suspected of being false matches). In many cases, especially if the gene of interest is well studied, the output of `searchForTerm()` will be entirely sufficient. Furthermore, due to the nature of organisation of metadata in SRA and GEO, `searchForTerm()` only conducts the tailored search within SRA (though it will add metadata from GEO for those samples that exist in both databases). Therefore, to ensure that no important samples have been missed, it might be beneficial to run:

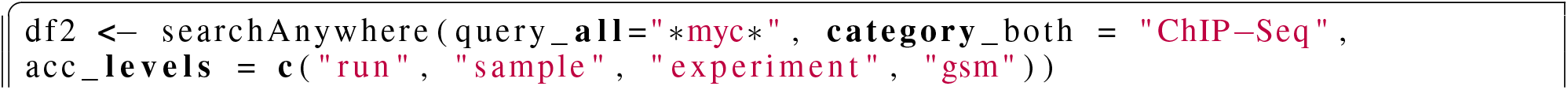
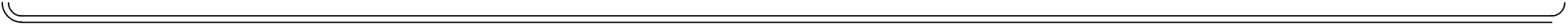

The `query_all` argument above denotes the query that will be used for both SRA and GEO databases (it is also possible to specify each separately in different arguments). `Category_both` is an optional argument referring to the study category (e.g. RNA-Seq, ChIP-Seq, WGS (Whole Genome Sequencing) etc.); GEO and SRA each use different naming systems (GEO refers to them as *type*, SRA uses *library strategy*), but `category_both` unifies them; to find out about the available options, please inspect the `SRA_GEO_Category_Conversion` object within the package. It is also possible to find out about the total number of entries within each category by running `getDatabaseInformation()`. It is also worth noting that a large number of studies in SRA are not classified, so it might desirable to either run a query without specifying the `category_both` argument, especially if searching for less common category types (e.g. Hi-C), as these might not always be classified. The last argument above, `acc_levels` determines which *accession levels* are taken into consideration for the search. For example, *study* level includes, among others, the study title and abstract, which contain more general information about the study; whereas the lower *accession levels* contain information about the samples themselves, such as details on treatment, cell lines etc.

It is worthwhile to check results with and without restriction to acc_levels. This can be done by either **(1)** running initial query with restriction (please note that by default the search is restricted, just like above; for further information please see function documentation) or **2)** including all accession levels and filtering out irrelevant results afterwards. Option **(2)** is likely preferable due to faster execution time.

Following the initial query, in case **(1)**, users may wish to search for remaining samples from the relevant studies, which might serve as ‘inputs’ for ChIP-Seq processing pipeline:

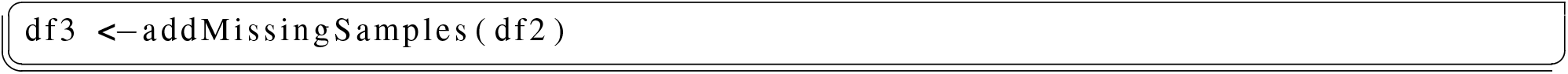

Approach **(2)**, might require further filtering of the data frame in order to refine results and only retrieve entries relevant to the user:

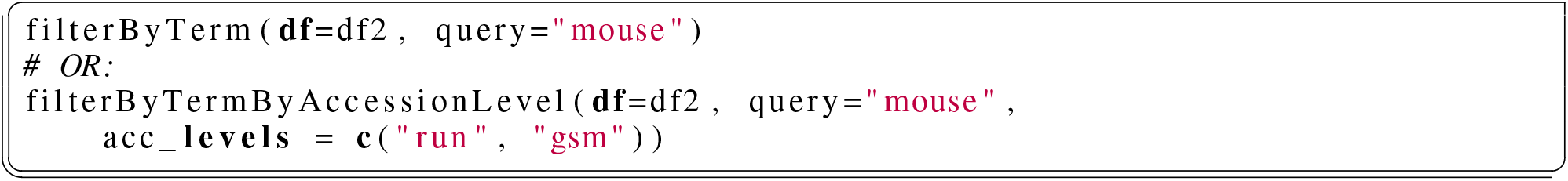

### 3.2 Further package usage examples

**’I would like to use SpiderSeqR to run queries’**

To run SpiderSeqR, users must initialise the environment by setting up the relevant database connections. This is done by one of the *setup functions*: `startSpiderSeqR()` and `startSpiderSeqRDemo()`

To setup a ‘demo’ version, which uses very small samples of the databases and allows users to get insights into the package functionality, please run the following at the beginning of each new session:

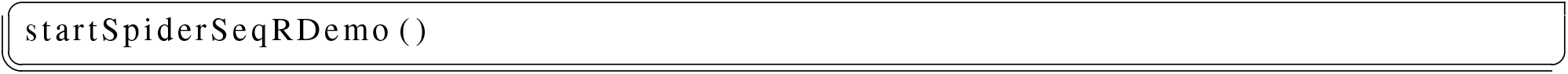

On the other hand, to use the real database files, please run:

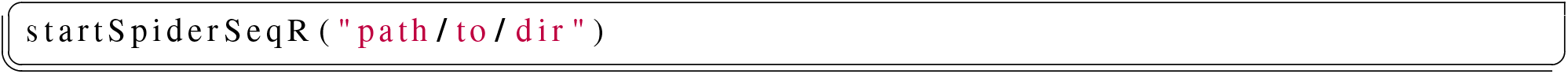

This function automatically downloads the database files (inside the specified directory), creates its own custom conversion database and sets up the database connections. Please allow time and disk space when running this function for the first time as the SRAdb and GEOmetadb files contain metadata about *millions* of samples (and they keep growing). However, once these files are in place, the function will only setup the connections, which takes a matter of seconds and should be repeated at the beginning of each new session. It is worth keeping the database files up to date; `startSpiderSeqR()` will prompt users to re-download the files if they become old; users can also choose the *expiry* arguments to either re-download the files earlier or to avoid the prompts to re-download.

**’I would like to find out more about the SRA(db) and GEO(metadb) databases’**

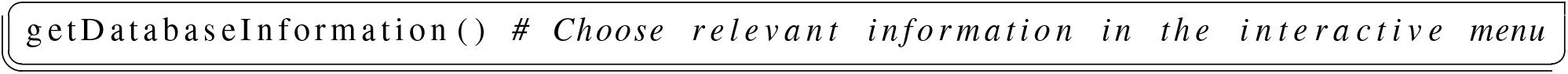

Users can gain insights into such aspects of the databases as number of entries (within each accession type), frequency counts for taxons and library strategies (e.g. RNA-Seq, ChIP-Seq, etc.). It is also possible to obtain a random sample from the database; all without the user having to use any SQLite queries.

**’I would like to rerun a query again, having updated the database files to discover new relevant data-sets’**

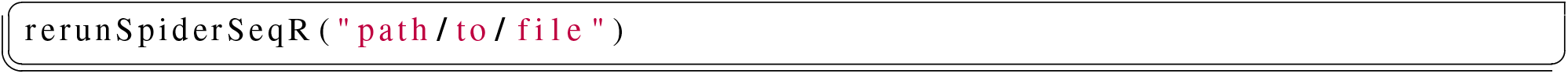

When running searches with any of the search functions (`searchForTerm()`, `searchAnywhere()`, `searchForAccession()`), users have the option to save the current query as a file. This enables one to easily re-run past queries; it might be most useful when periodically checking for new experiments matching a complex query run in the past.

## 4 DISCUSSION

Scientific research is a costly endeavour, and is mostly funded by taxpayers or charities. The ability to reuse data to advance research and innovation, reproduce studies by re-anlysing data to validate, verify and confirm previous research, and to make publicly funded data-assets available to the public, thereby leveraging investment into research are some of the key benefits of open access research data [14]. These practices also lead to utilising shared data-sets to produce new knowledge [15] and potential to implement FAIR data principles making data findable, accessible, inter-operable and reusable [16].

SRA and GEO are truly useful projects, enabling sharing and exchange of ‘omic’ data, which is important for data reuse [17]. An example demonstrates reuse of large amounts of data (over 5000 gene expression data-sets from GEO), where the different studies were integrated and revealed the overall structure of gene expression space across different tissues, which could not have been observed using any of the contributing studies individually [18]. Similarly, Tsui et al., reprocessed over 250,000 human sequencing data-sets (more than 1000 TB data worth of raw sequence data) from SRA into a single unified data-set of allelic read counts for nearly 300,000 variants of biomedical relevance demonstrating the usefulness of data reuse [19]. However, there are some significant obstacles, some inherent to the database design itself, some related to metadata quality, which hinders efficient reuse of the data.

In order to more efficiently re-purpose sequencing data-sets in multi-omic data repositories we built the R package SpiderSeqR. Currently it provides search and filtering access to data and metadata in NCBI GEO and SRA databases. While there are a number of other tools and resources to access these publicly available sequencing data-sets, most of these have limitations. A comparison of the available tools, in Table 1 will enable users select an option best suited to their needs. The simplest means of searching the NCBI SRA and GEO databases are through their web interfaces [3, 7]. These web interfaces can be used to search for a specific accession or search a term with a handful of matching experiments. However, searching and retrieving large data-sets or filtering through a few hundred search results can be time consuming and difficult.

**Table 1:**
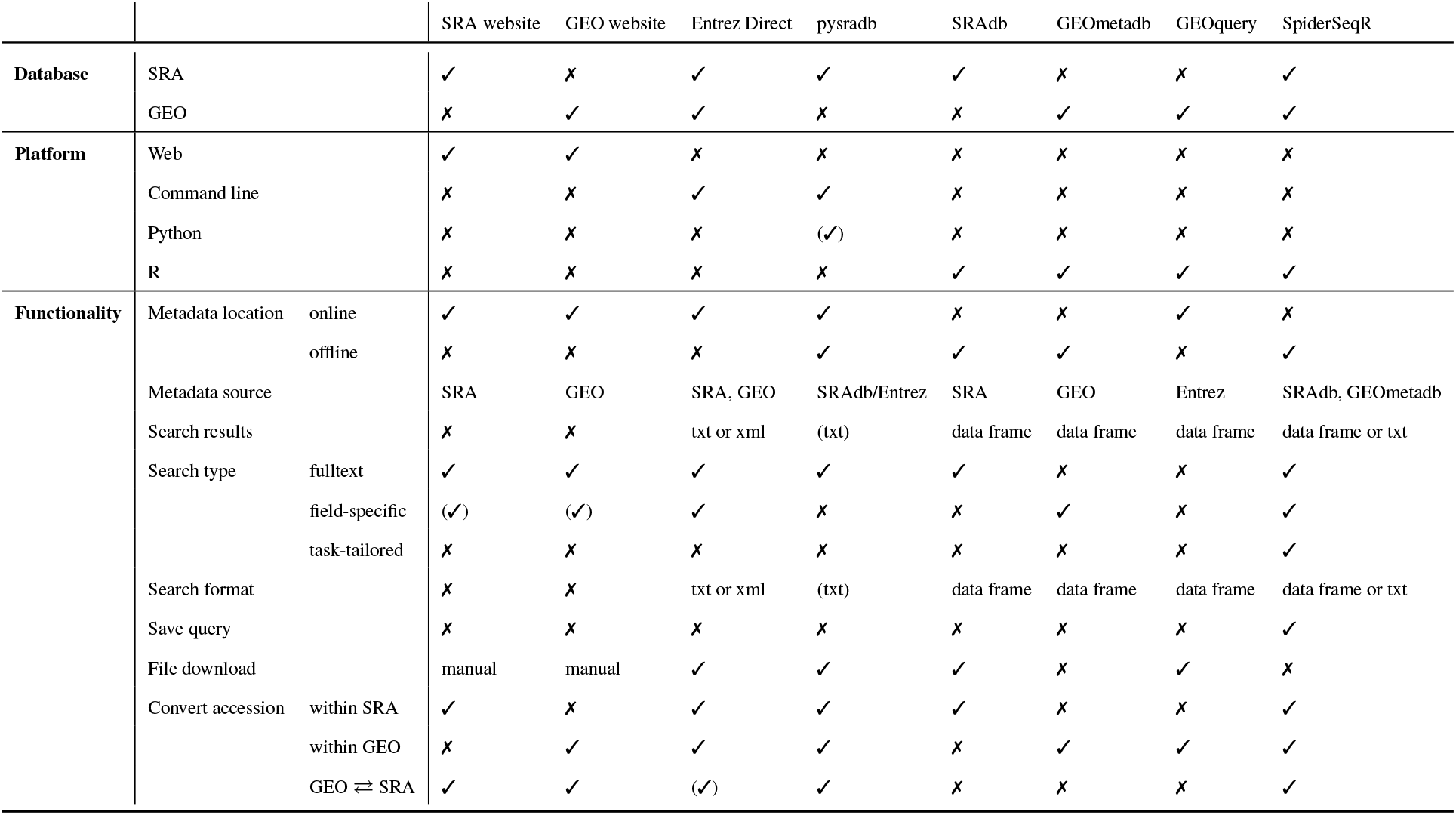
Comparison of the tools for search and retrieval of data from SRA and GEO databases.

The next available option is to use Entrez command line tools [20]. The Entrez Programming Utilities (E-utilities) are a set of nine server-side programs that provide a stable interface into the Entrez query and database system at the NCBI. The E-utilities use a fixed URL syntax that translates a standard set of input parameters into the values necessary for various NCBI software components to search for and retrieve the requested data. The E-utilities are therefore the structured interface to the Entrez system, which currently includes 38 databases covering a variety of biomedical data, including nucleotide and protein sequences, gene records, three-dimensional molecular structures, and the biomedical literature. The advantage of this tool is that it can be accessed programmatically, but there are no further tools available for filtering or refining the results. Although ample documentation is available about Entrez E-Utils, their usage is not intuitive.

Another option is SRAdb [21] and GEOmetadb [22], two Bioconductor/R packages. They allow for searching the databases using the R programming language, which makes the results more convenient to manipulate and integrate into workflows. However, the search capabilities are fairly simple (SQLite queries and full-text search for SRAdb and SQLite only for GEOmetadb) and there is no integration between the two databases - each of the two packages acts as a separate entity, so to search all the samples, one has to perform individual queries within SRAdb and GEOmetadb. The GEOquery package is yet another Bioconductor interface to NCBI GEO [23]. Probably the most versatile tool to date is pysradb [24], which offers a degree of integration between SRA and GEO. It is a command line tool written in Python. However, because it is command line based, it makes further refinements more difficult. Disappointingly, it also offers only a limited amount of information about the samples, compared to that available online or in the parsed SRAdb/GEOmetadb databases.

SpiderSeqR aims to provide both functionalities missing among the existing tools and a simple to use intuitive toolkit. Firstly, it has two main search functions; searchAnywhere (fulltext) and searchForTerm (tailored to minimise spurious matches). The latter can additionally label suggested *inputs* and *controls* in ChIP-Seq and RNA-Seq experiments respectively. Furthermore, we provide convenient tools for filtering the results. There are also functions allowing for saving the record of the search query, meaning that it can be reused in the future (e.g. a complicated and well refined query can be run periodically on an updated version of the database, to check for any new experiments). Importantly, SpiderSeqR allows for seamless integration between GEO and SRA, meaning that it is not necessary to search both individually. It’s modular nature and foundations based on SQLite data structures will enable future implementations of other biomedical databases and data types.

As mentioned before, both GEO and SRA are organised by a hierarchy of accessions. The top level in GEO is the *series* record that provides a focal point and description of the whole study and is assigned a unique and stable GSE accession identifier. However there is also a higher level accession, known as *superseries* (and is also assigned a GSE accession). Confusingly, it is impossible to distinguish between series and superseries based on the accession number itself. Not all samples belong to superseries, but those that do, have minimum two GSEs assigned to them (i.e. series and superseries one), which makes it difficult to appreciate the relationships between samples within series and to search for other samples from the same series. SpiderSeqR mitigates this problem by searching within all GSEs that any of the samples belongs to. Within a series are sample accessions (GSM). A sample record describes conditions under which individual samples were handled, including the manipulation it underwent. Sample entities must reference only one Platform and may be included in multiple series. Similarly, in SRA the top level record is the project identified using an SRP accession. Usually these are equivalent to an GSE accession in GEO. SRX contains experiment level data and SRR contains individual runs.

The metadata of SRA and GEO databases are generated by the submitting researchers, leading to a large degree of variability in the style and quality of sample descriptions. The databases have guidelines regarding required information within each field, but there is a wide range of possible approaches one could use when making a submission, leading to missing experiment details, steps and reagent annotation. These disparities are very difficult to overcome programmatically (or perhaps even with natural language processing solutions). The only definitive solution would be for SRA and GEO to introduce metadata quality checks at the point of submission and further measures to promote more uniform quality of metadata, allowing for more efficient reuse of data in the future.

The functionality of SpiderSeqR can be easily extended for data download by installing the SRA Toolkit https://ncbi.github.io/sra-tools/. Similarly, it is also possible to download GEO data using the Entrez utilities. This will enable direct download of data-sets using the sample sheets containing filtered search results generated by SpiderSeqR.

In summary, SpiderSeqR is an R package providing integration between SRA and GEO databases, handling the accession conversions, full-text, SQLite and tailored search, as well as control of display and filtering/refining the results to allow for more efficient search of the two databases.

## AUTHOR CONTRIBUTIONS

S.A.S conceived the study. A.M.S developed the methods and carried out the software engineering, C.F was involved in the initial stages during a summer studentship, D.B helped with testing and analysis. A.M.S and S.A.S wrote the manuscript with input from the other authors.

## ACKNOWLEDGEMENTS

A.M.S, C.F, D.B and S.A.S were supported by the UK Medical Research Council core funding (MC_UU_12022/10) to the S.A.S laboratory. We also thank members of the S.A.S laboratory that read and commented on the manuscript.

